# Loss of Dna2 nuclease activity results in decreased Exo1-mediated resection at DNA double strand breaks

**DOI:** 10.1101/2023.10.25.564088

**Authors:** Aditya Mojumdar, Courtney Granger, Martine Lunke, Jennifer A. Cobb

## Abstract

A DNA double strand break (DSB) is one of the most dangerous types of DNA damage that is repaired largely by homologous recombination (HR) or non-homologous end-joining (NHEJ). The interplay of repair factors at the break directs which pathway is used, and a subset of these factors also function in more mutagenic alternative (alt) repair pathways. Resection is a key event in repair pathway choice and extensive resection, which is a hallmark of HR, is mediated by two nucleases, Exo1 and Dna2. We observed differences in resection and repair outcomes in cells harbouring nuclease dead *dna2*-1 compared to *dna2*Δ *pif1*-m2 that could be attributed to the level of Exo1 recovered at DSBs. Cells harbouring *dna2*-1 showed reduced Exo1 localization, increased NHEJ, and a greater defect in resection compared to cells where *DNA2* was deleted. Both the decreased level of resection and the increased rate of NHEJ in *dna2*-1 mutants were reversed upon deletion of *KU70* or ectopic expression of Exo1. By contrast, when *DNA2* was deleted, Exo1 and Ku70 recovery levels did not change, however Nej1 increased as did the frequency of alt-EJ/ MMEJ repair. Our findings demonstrate that decreased Exo1 at DSBs contributed to the resection defect in cells expressing inactive Dna2 and highlight the complexity of understanding how functionally redundant factors are regulated *in vivo* to promote genome stability.

**Graphical Abstract:** 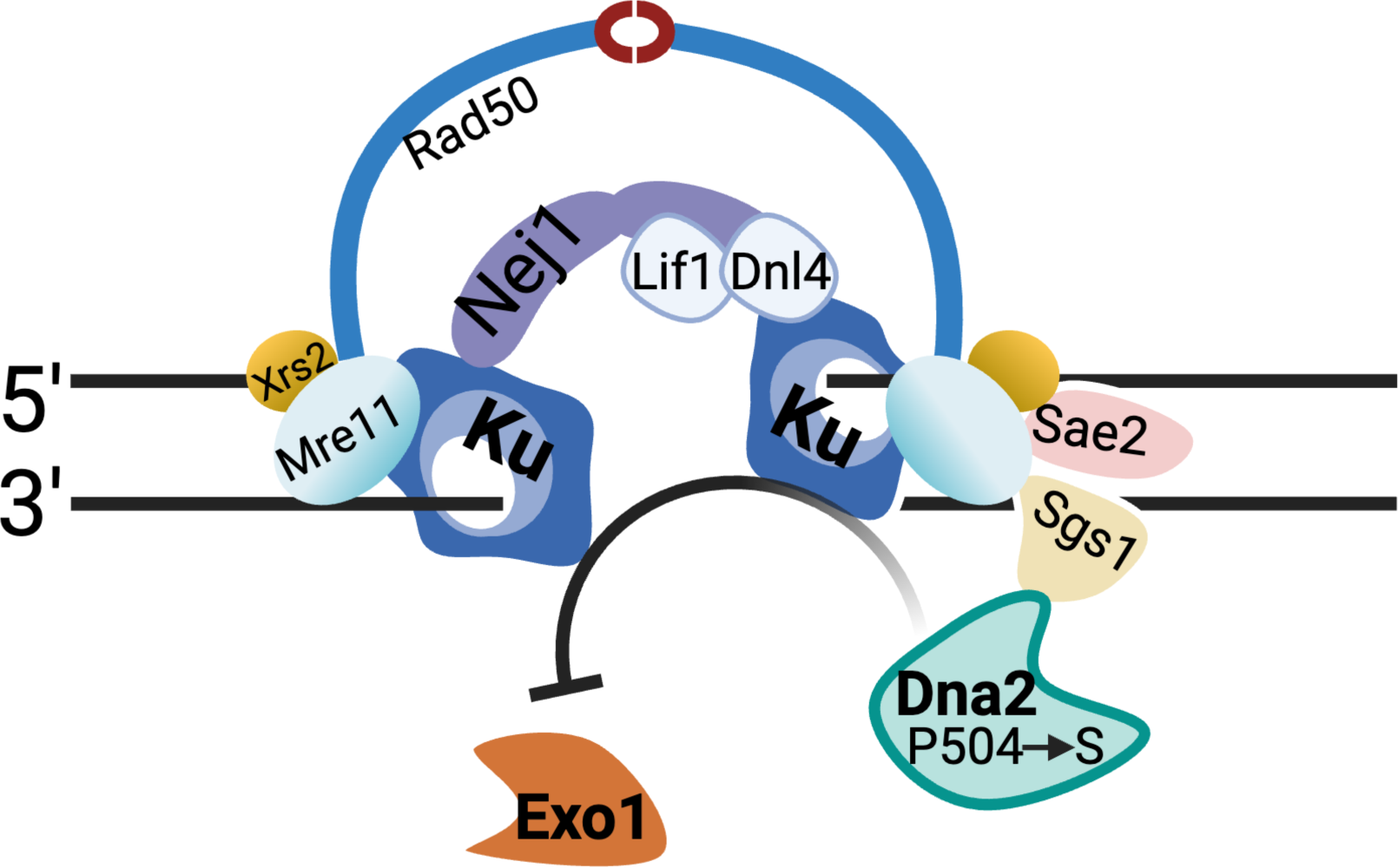

## Introduction

Homologous recombination (HR) and non-homologous end joining (NHEJ) are the canonical pathways of DNA double-strand break (DSB) repair. HR is an error-free pathway requiring extensive 5’ end resection and NHEJ is an error-prone pathway whereby the ends are joined after minimal processing. DNA resection is the major deciding step between these two pathways [1]. However, if resection initiates and HR is not possible, then a more mutagenic alternative (alt) repair pathway can be used as a last resort. Microhomology-mediated end joining (MMEJ) is an alt-end-joining (EJ) pathway that occurs at a high frequency in the absence of Ku or when broken ends are not compatible for direct ligation. MMEJ requires 5’ resection, however in contrast to HR, the extent of resection in MMEJ is believed to be coordinated with the process of scanning for microhomology (MH) in adjacent regions flanking the DSB. The mechanism remains ill-defined, however the repair product from MMEJ contains a deletion corresponding in size to the fragment between the annealed MH sequences, which were revealed during resection.

The first responders to a DSB are yKu70/80 (Ku) and Mre11-Rad50-Xrs2 (MRX) and they are important for recruiting additional NHEJ and HR factors [2–6]. The Ku heterodimer also protects the ends from nucleolytic degradation and aids in the localization of Lif1-Dnl4 and Nej1 [3]. Dnl4 ligase completes end-joining by ligating the DNA with the help of Lif1 and Nej1 [4, 7–9]. The central role of the MRX complex is to tether the loose DNA ends mainly through the structural features of Rad50 [10, 11] and to initiate resection through the nuclease activity of Mre11 [12]. Sae2 interacts with the MRX complex and activates Mre11 nuclease activity, which forces Ku dissociation. Ku disengagement at the DSB coincides with the initiation of 5’ to 3’ end-resection by two long-range resection nucleases, Dna2 in complex with Sgs1 helicase, and Exo1 [12–14]. Dna2 and Exo1 nucleases show functional redundancy as Exo1 drives long-range resection in the absence of Dna2, and vice versa [14]. The interplay between repair factors in the two canonical pathways regulate the initiation of resection in part through antagonistic relationships between Ku and Exo1, and between Nej1 and Dna2, wherein Nej1 inhibit interactions of Dna2 with Sgs1 and with Mre11 and Sae2 [5, 6, 15, 16].

Dna2 is mutated in a myriad of human cancers, but due to its essential role in processing Okazaki fragments and other intermediates at replication forks, *DNA2* cannot be deleted [17–21]. However, in yeast *dna2*Δ lethality can be supressed by mutation of *PIF1,* a gene encoding a DNA helicase that does not function in 5’ resection at DSBs [22]. In the absence Mre11 nuclease activity, resection initiates primarily through Dna2, not through Exo1 [23–25]. However, there is a gap in understanding the regulation of Dna2 at DSBs as nuclease deficient *dna2*-1 (P504S) shows greater sensitivity to DSB-inducing agents compared to dna2Δ *pif1*-m2 (Fig. 1A) [26, 27].

**Figure 1.**
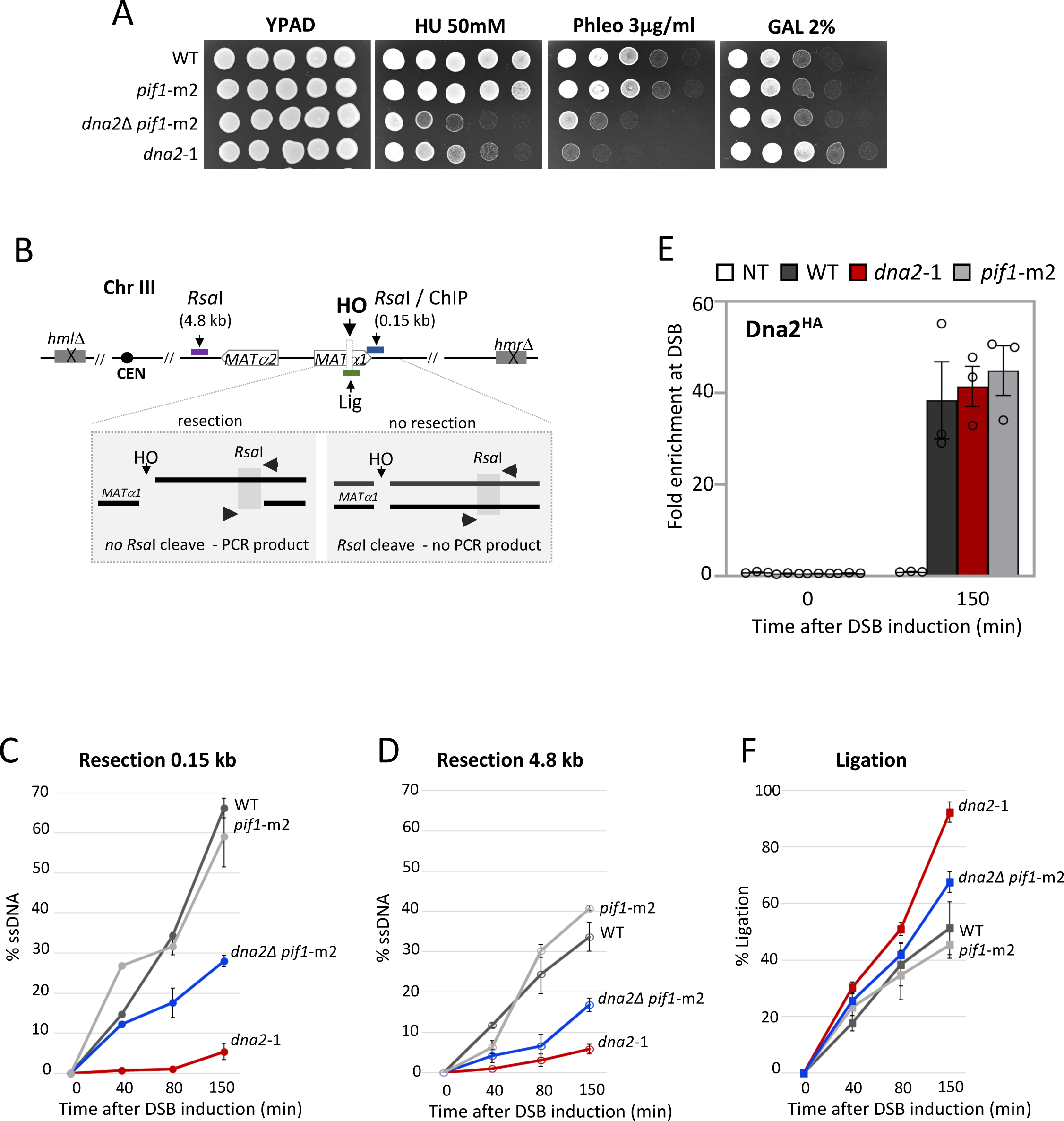
Nuclease deficient *dna2*-1 leads to compromised resection at DSB. **(A)** Five-fold serial dilutions of the strains - wild type (JC-727), *dna2*Δ *pif1*-m2 (JC-6005), *pif1*-m2 (JC-6006) and *dna2*-1 (JC-6007) were spotted on YPAD, 50mM HU, 3.0 μg/ml phleomycin and 2% galactose containing plates. **(B)** Schematic representation of regions around the HO cut site on chromosome III. The ChIP primers used in this study correspond to 0.15kb (blue) from the DSB and the end-joining primers flank the HO cut site (Lig, green). The qPCR resection assay relies on two *Rsa*I sites located 0.15 kb (blue), and 4.8 kb (purple)from the DSB. **(C-D)** qPCR-based resection assay of DNA 0.15 kb and 4.8 kb away from the HO DSB, as measured by % ssDNA, at 0, 40, 80 and 150 mins post DSB induction in cycling cells in wild type (JC-727), *dna2*Δ *pif1*-m2 (JC-6005), *pif1*-m2 (JC-6006) and *dna2*-1 (JC-6007). **(E)** Enrichment of Dna2^HA^ at 0.15kb from DSB 0 min (no DSB induction) and 150 mins after DSB induction in wild type (JC-4117), *dna2*-1 (JC-5707), *pif1*-m2 (JC-6130) and no tag control (JC-727) was determined. The fold enrichment is normalized to recovery at the *PRE1* locus. **(F)** qPCR-based ligation assay of DNA at HO DSB, as measured by % Ligation, at 0, 40, 80 and 150 mins in cycling cells in glucose post DSB. Strains used were wild type (JC-727), *dna2*Δ *pif1*-m2 (JC-6005), *pif1*-m2 (JC-6006) and *dna2*-1 (JC-6007).

Here we determined that the dominant negative effects of *dna2*-1 were caused by decreased localization of Exo1 nuclease, the factor functionally redundant with Dna2 in DSB repair. In *dna2*-1 mutant cells, Ku-dependent NHEJ increased and Exo1-dependent 5’ resection decreased. By contrast, in dna2Δ *pif1*-m2 mutants, Ku70 and Exo1 recruitment to the break remained indistinguishable from wild type, but end-joining repair occurred mainly through MMEJ. These results demonstrate that Exo1 is impacted by the physical presence of Dna2 and that the interplay between these two nucleases regulates key events that drive repair pathway choice, including the ratio of NHEJ and MMEJ.

## Results

### Nuclease deficient dna2-1 show abrogated resection at DSB

Cells harbouring nuclease dead *dna2*-1 were more sensitive than *dna2Δ pif1*-m2 mutants to Phleomycin, an agent that causes DNA double strand breaks, but less sensitive to hydroxyurea (HU), an agent inducing replication stress (Fig. 1A, [26, 27]). While sensitivities to various genotoxic stressors have been previously reported with *dna2* mutants, there has been little work explaining why *dna2*-1 mutants show greater sensitivity to DSB causing agents compared to cells harbouring the deletion of *DNA2* or its binding partner, *SGS1* (Figs. 1A and S1). This prompted our side-by-side investigation of *dna2Δ pif1*-m2 and *dna2*-1 in DSB repair. Our aims were to evaluate DNA resection, a key early step in HR and then to determine the impact of the *dna2* mutations on the functionality of the other DSB repair factors.

Dna2 functions in long-range resection and can compensate for Mre11 to initiate resection [14]. To this end, resection was determined at two locations, 0.15 and 4.8 kb from the HO-induced DSB using a quantitative PCR-based approach that relies on *Rsa*I as previously described [16, 30]. Resection produces single-stranded DNA (ssDNA) and if resection goes beyond the *Rsa*I recognition sequence, then the site is not cleaved and can be amplified by PCR (Fig. 1B). Resection at the time points (0-150 mins) was similar at both distances from the break in *pif1*-m2 and wild type, indicating that the loss of *PIF1* activity did not impact DNA processing at the DSB (Figs. 1C, D). Furthermore, when we performed chromatin immuno-precipitation (ChIP) at the HO-induced DSB, Dna2^HA^ levels in *pif1*-m2 mutants were indistinguishable from wild type (Fig. 1E), reinforcing earlier work showing that the disruption of *PIF1* did not impact DSB repair [22]. Upon deletion of *DNA2*, resection decreased by ∼2-fold, with a slightly greater defect at the distance 4.8 kb from the break (Figs. 1C, D). A more pronounced defect was observed in *dna2*-1 mutants as resection was abrogated at both distances (Figs. 1C, D). Dna2 recovery in *dna2*-1 mutants was unaltered (Fig. 1E), suggesting the physical association of nuclease-dead Dna2 at the break had a dominant negative impact.

We also observed that *dna2*-1 mutant cells survived better than *dna2Δ pif1*-m2 and WT on 2% GAL (Fig. 1A). The genetic background of these cells includes *hml*11 *hmr*11 which prevents HR. Survival on galactose therefore correlates with mutagenic end-joining repair, which prevents subsequent HO-cutting as opposed to survival on Phleomycin, which creates multiple DSBs throughout the genome that can repair by HR.

To complement the survival assays, we performed end-joining experiments where the DSB was induced with galactose for 2 hours before cells were wash and released into glucose to prevent further re-cutting. At the indicated time points, genomic DNA was prepared, and quantitative PCR was performed with primers spanning the HO recognition site as previously described [9]. The rate of end-joining increased more in *dna2*-1 compared to *dna2Δ pif1*-m2 mutants (Fig. 1F). Increased end-joining might arise naturally as a consequence of decreased HR, but could also arise from more NHEJ factors at the DSB in *dna2*-1 mutants.

### NHEJ factors at DSBs in dna2 mutants

Prior to comparing the impact of the *dna2* mutants on factors driving resection, we determined the localization of proteins essential for NHEJ. Ku70 recovery at the DSB increased in *dna2*-1 mutant cells but not in *dna2*Δ *pif1*-m2 mutants (Fig. 2A). By contrast, the recovery of Nej1 increased significantly in both mutants, with *dna2*Δ *pif1*-m2 showing a greater increase (Fig. 2B). These data highlight the antagonistic relationship between Dna2 and Nej1 at DSBs [5]. The recovery of the other canonical NHEJ factors Lif1 and Dnl4, in cells harbouring either of the *dna2* mutant were indistinguishable from wild type (Figs. S2A,B).

**Figure 2.**
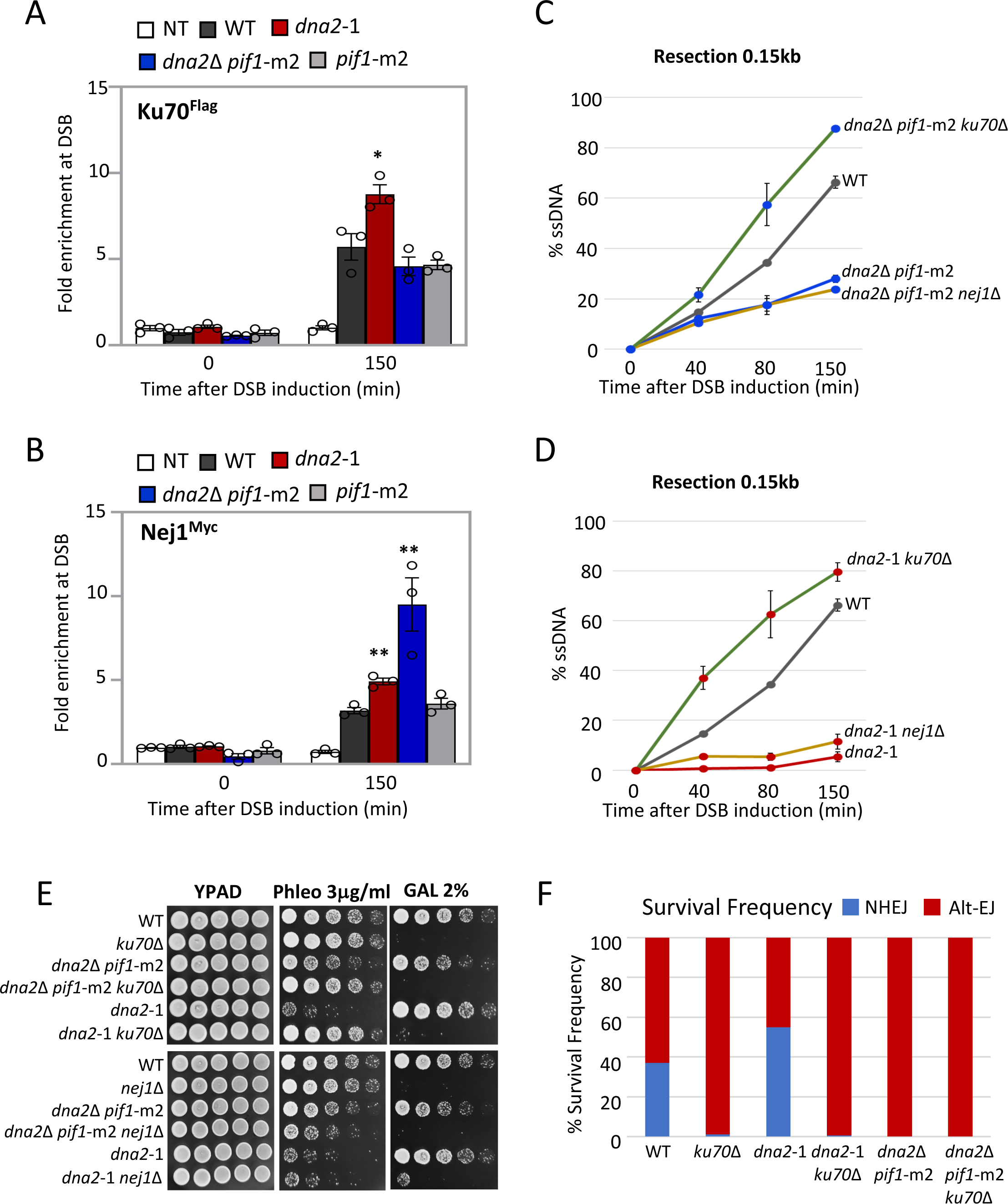
Nuclease deficient *dna2*-1 promotes end-joining at DSB. **(A)** Enrichment of Ku70^Flag^ at 0.15kb from DSB, 0 min (no DSB induction) and 150 mins after DSB induction in wild type (JC-3964), *dna2*-1 (JC-6237), *dna2*Δ *pif1*-m2 (JC-6068), *pif1*-m2 (JC-6069) and no tag control (JC-727) was determined. The fold enrichment is normalized to recovery at the *PRE1* locus. **(B)** Enrichment of Nej1^Myc^ at 0.15kb from DSB, 0 min (no DSB induction) and 150 mins after DSB induction in wild type (JC-1687), *dna2*-1 (JC-5479), *dna2*Δ *pif1*-m2 (JC-6099), *pif1*-m2 (JC-6132) and no tag control (JC-727). **(C-D)** qPCR-based resection assay of DNA 0.15 kb away from the HO-DSB, as measured by % ssDNA, at 0, 40, 80 and 150 mins post DSB induction in cycling cells in wild type (JC-727), *dna2*Δ *pif1*-m2 (JC-6005), *ku70*Δ *dna2*Δ *pif1*-m2 (JC-6128), *nej1*Δ *dna2*Δ *pif1*-m2 (JC-6060), *dna2*-1 (JC-6007), *ku70*Δ *dna2*-1 (JC-5942) and *nej1*Δ *dna2*-1 (JC-5670). **(E)** Five-fold serial dilutions of the strains used in (C) and (D) were spotted on YPAD, 3.0 μg/ml phleomycin and 2% galactose containing plates. **(F)** Survival frequencies depicting the ratio of NHEJ (blue) and alt-EJ/MMEJ (red) repair frequencies in wild type (JC-5903), *ku70*Δ (JC-6195), *dna2*-1 (JC-6105), *ku70*Δ *dna2*-1 (JC-6273), *dna2*Δ *pif1*-m2 (JC-6181) and *ku70*Δ *dna2*Δ *pif1*-m2 (JC-6280). For all the experiments - error bars represent the standard error of three replicates. Significance was determined using a 1-tailed, unpaired Student’s t test. All strains compared are marked (P<0.05*; P<0.01**) and compared to WT.

We wanted to determine whether preventing NHEJ would reverse the resection defect of the *dna2* mutants, and potentially the dominant negative effect of *dna2*-1 in HR repair. In addition to their essential function in NHEJ, both Ku70 and Nej1 inhibit resection (Fig. S2E). In alignment with our earlier work, deletion of *KU70*, but not the other core NHEJ factors, reversed the resection defect in both *dna2* mutants (Figs. 2C-D and S2C-D; [9]). Correlating with the rescue in resection, deletion of *KU70*, but not *NEJ1*, suppressed the phleomycin sensitivities of both *dna2*Δ *pif1*-m2 and *dna2*-1 mutants (Fig. 2E). These data indicate that suppressing the dominant negative resection defect of *dna2*-1 was specific to the loss of Ku70 rather than disruption of NHEJ by *ku70*Δ as HR-mediated repair was restored in *dna2*-1 *ku70*Δ, but not in *dna2*-1 *nej1*Δ mutants.

To determine the type of end-joining that can proceed in these mutants we utilized a reporter system containing a *URA3* marker flanked by two inverted HO recognition sites (Fig. S2F) [31]. If both sites are cleaved simultaneously, non-compatible ends are generated and alt-EJ/ MMEJ is used. However, because cutting at both sites is not perfectly coordinated, each single cut can still be repaired by NHEJ as previously described [31]. In wild type cells, the relative frequency of NHEJ and alt-EJ/MMEJ as determined by growth on -URA was 37% and 63% respectively (Fig. 2F). The relative frequencies of NHEJ and MMEJ notably differed between the two *dna2* mutants. The frequency of NHEJ increased to 55% in *dna2*-1 mutants (Fig. 2F). By contrast, when *DNA2* was deleted, alt-EJ/MMEJ was the preferred end joining pathway, which we found surprising given Ku70 was similarly recovered at the break site in *dna2*Δ *pif1*-m2 and wild type cells (Figs 2A, F). Consistent with previous work, upon *KU70* deletion end joining occurred through alt-EJ/MMEJ, and NHEJ in *dna2*-1 mutants was dependent on Ku70 (Fig. 2F).

### Nuclease deficient dna2-1 suppresses Exo1 recruitment to DSB

To bring further insight to events underlying the *dna2*-1 phenotype, we also determined the impact of *dna2*-1 and *DNA2* deletion on the localization of other factors important for resection, namely Exo1 nuclease and the nuclease-associated factors, Sae2 and Sgs1. The recovery of Sgs1 and Sae2 were not altered in either mutant background (Fig. S3A, B). By contrast, Exo1 recovery was significantly reduced in *dna2*-1 to almost the level of the non-tagged (NT) control whereas its recovery in *dna2*Δ *pif1*-m2 was like wild type (Fig. 3A). These data indicate that localization of Exo1 was inhibited by the presence of nuclease deficient Dna2 at the break rather than the intrinsic loss of Dna2 nuclease activity.

**Figure 3.**
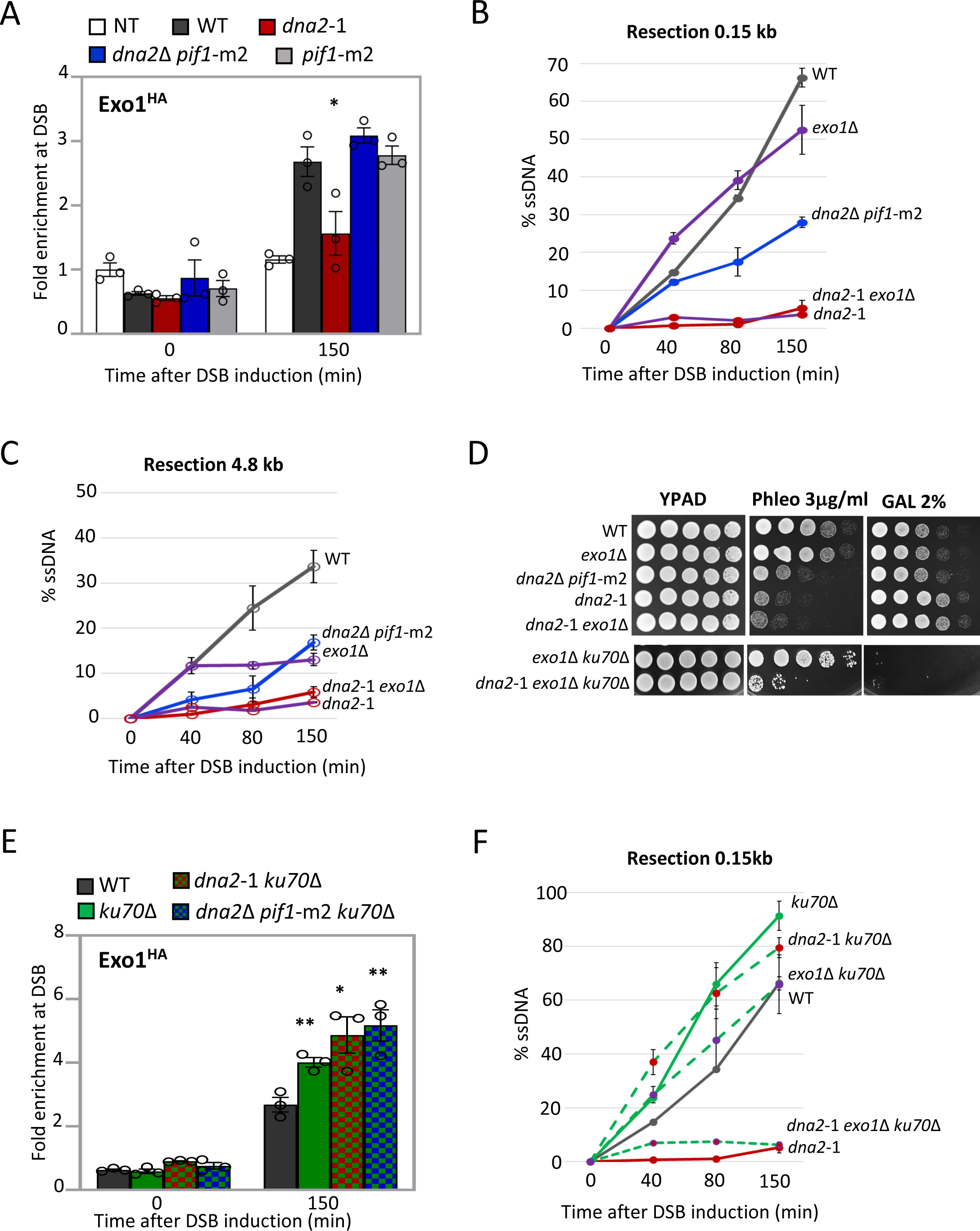
Nuclease deficient *dna2*-1 suppresses Exo1 recruitment at DSB. **(A)** Enrichment of Exo1^HA^ at 0.15 kb from DSB 0 min (no DSB induction) and 150 mins after DSB induction in wild type (JC-4869), *dna2*-1 (JC-6020), *dna2*Δ *pif1*-m2 (JC-6115), *pif1*-m2 (JC-6117) and no tag control (JC-727) was determined. The fold enrichment is normalized to recovery at the *PRE1* locus. **(B-C)** qPCR-based resection assay of DNA 0.15 kb and 4.8 kb away from the HO DSB, as measured by % ssDNA, at 0, 40, 80 and 150 mins post DSB induction in cycling cells in wild type (JC-727), *exo1*Δ (JC-3767), *dna2*-1 (JC-6007), *exo1*Δ *dna2*-1 (JC-5692) and *dna2*Δ *pif1*-m2 (JC-6005). **(D)** Five-fold serial dilutions of wild type (JC-727), *exo1*Δ (JC-3767), *dna2*-1 (JC-6007), *exo1*Δ *dna2*-1 (JC-5692), *dna2*Δ *pif1*-m2 (JC-6005), *ku70*Δ *exo1*Δ (JC-3837) and *ku70*Δ *exo1*Δ *dna2*-1 (JC-6025) were spotted on YPAD, 3.0 μg/ml phleomycin and 2% galactose containing plates. **(E)** Enrichment of Exo1^HA^ at 0.15 kb from DSB, 0 min (no DSB induction) and 150 mins after DSB induction in wild type (JC-4869), *ku70*Δ (JC-6018), *ku70*Δ *dna2*-1 (JC-6215) and *ku70*Δ *dna2*Δ *pif1*-m2 (JC-6213) was determined. The fold enrichment is normalized to recovery at the *PRE1* locus. **(F)** qPCR-based resection assay of DNA 0.15kb away from the HO DSB, as measured by % ssDNA, at 0, 40, 80 and 150 min post DSB induction in cycling cells in wild type (JC-727), *ku70*Δ (JC-1904), *dna2*-1 (JC-6007), *ku70*Δ *dna2*-1 (JC-5942), *ku70*Δ *exo1*Δ (JC-3837) and *ku70*Δ *exo1*Δ *dna2*-1 (JC-6025). For all the experiments - error bars represent the standard error of three replicates. Significance was determined using a 1-tailed, unpaired Student’s t test. All strains compared are marked (P<0.05*; P<0.01**) and compared to WT.

We next compared resection and survival when the *dna2* mutants were combined with deletion of *EXO1*. In line with previous observations, short-range resection (0.15 kb) decreased modestly in *exo1*Δ single mutant cells (Fig. 3B), but there was a ∼ 2-fold decrease in long-range resection (4.8 kb) 80-150 mins after DSB induction (Fig. 3C) [14, 16]. We could not determine resection when both nucleases were deleted due to synthetic lethality (SL) [14]. However, resection at both distances from the break and phleomycin sensitivity in *dna2*-1 *exo1*Δ mutants was indistinguishable from *dna2*-1 (Figs. 3B-D). One explanation for the marked decrease in resection in *dna2*-1 single mutants was that nuclease deficient Dna2 directly blocked the association of Exo1 with DSBs. However, Exo1 was similarly recovered in *DNA2*+ wild type cells and *dna2*Δ *pif1*-m2 mutant cells, thus the presence or absence of Dna2 per se did not directly impact Exo1 recovery. A more plausible model, based on previous work showing Exo1 to be negatively regulated by the presence of Ku [15], is that the pronounced resection defect in *dna2*-1 mutants resulted from decreased Exo1 because of increased Ku, in addition to the loss of Dna2 nuclease activity.

Indeed, upon deletion of *KU70*, Exo1 recovery increased at the DSB in *dna2*-1 mutants (Fig 3E). These data were also consistent with suppression of *dna2*-1 phleomycin sensitivity by *KU70* deletion being Exo1 dependent (Figs. 2D and 3D). Resection remained low in *dna2*-1 *exo1*Δ *ku70*Δ triple mutants, as did growth on phleomycin and 2% GAL because both main DSB repair pathways, HR and NHEJ, were disrupted (Fig. 3D, F). However, alt-EJ /MMEJ was still functional, which could provide some insight as to how a small percentage of triple mutants survived on phleomycin (Figs. 3D and S3C). Lastly, Exo1 recovery also increased when *KU70* was deleted in *dna2*Δ *pif1*-m2 (Fig 3E), which helps to explain why the resection defect in *dna2*Δ *pif1*-m2 mutants was supressed by *ku70*Δ, but not by *nej1*Δ (Fig. 2C). Our earlier work showed the inhibitory effect of Nej1 on resection to be unrelated to Exo1 activity [5, 6, 9].

### Overexpression of Exo1 restores the resection defect in dna2-1 cells

Our data thus far support a model whereby the dominant negative affect of *dna2*-1 on resection stemmed from increased Ku70 at the DSB, which in turn inhibited Exo1 localization. This was supported by genetic analysis where deletion of *KU70* in *dna2*-1 resulted in increased Exo1 recovery, increased resection, and decreased sensitivity to phleomycin. We next wanted to determine whether increasing the level of Exo1 could rescue the dominant negative effect of *dna2*-1 mutants. We utilized a 2μ URA3 plasmid encoding Exo1 (pEM-EXO1) that was previously engineered to investigate the inhibition Exo1 by Ku [15]. Of note, expression of Exo1 did not alter Dna2 recovery in wild type or *dna2*-1 mutants (Fig. 4A). Resection at 0.15kb in *dna2*-1 + pEM-EXO1 increased to the level seen in *dna2*Δ *pif1*-m2 mutants, although resection in both remained lower than wild type (Fig. 4B). At the further distance, 4.8kb from the DSB, Exo1 expression in both *dna2* mutants resulted in a greater rescue where resection in *dna2*-1 + pEM-EXO1 was like wild type + EV, and resection in both *dna2*Δ *pif1*-m2 and wild type + pEM-EXO1 were similarly increased (Fig. 4C). Highlighting the link between resection and *in vivo* DSB repair, Phleomycin sensitivity decreased in both *dna2* mutants expressing Exo1 most notably in *dna2*-1 mutants after 3 days of growth (Fig. 4D). Resection and phleomycin sensitivity in *pif1*-m2 + pEM-EXO1 was indistinguishable from wild type (Figs. 4D and S4A, B)

**Figure 4.**
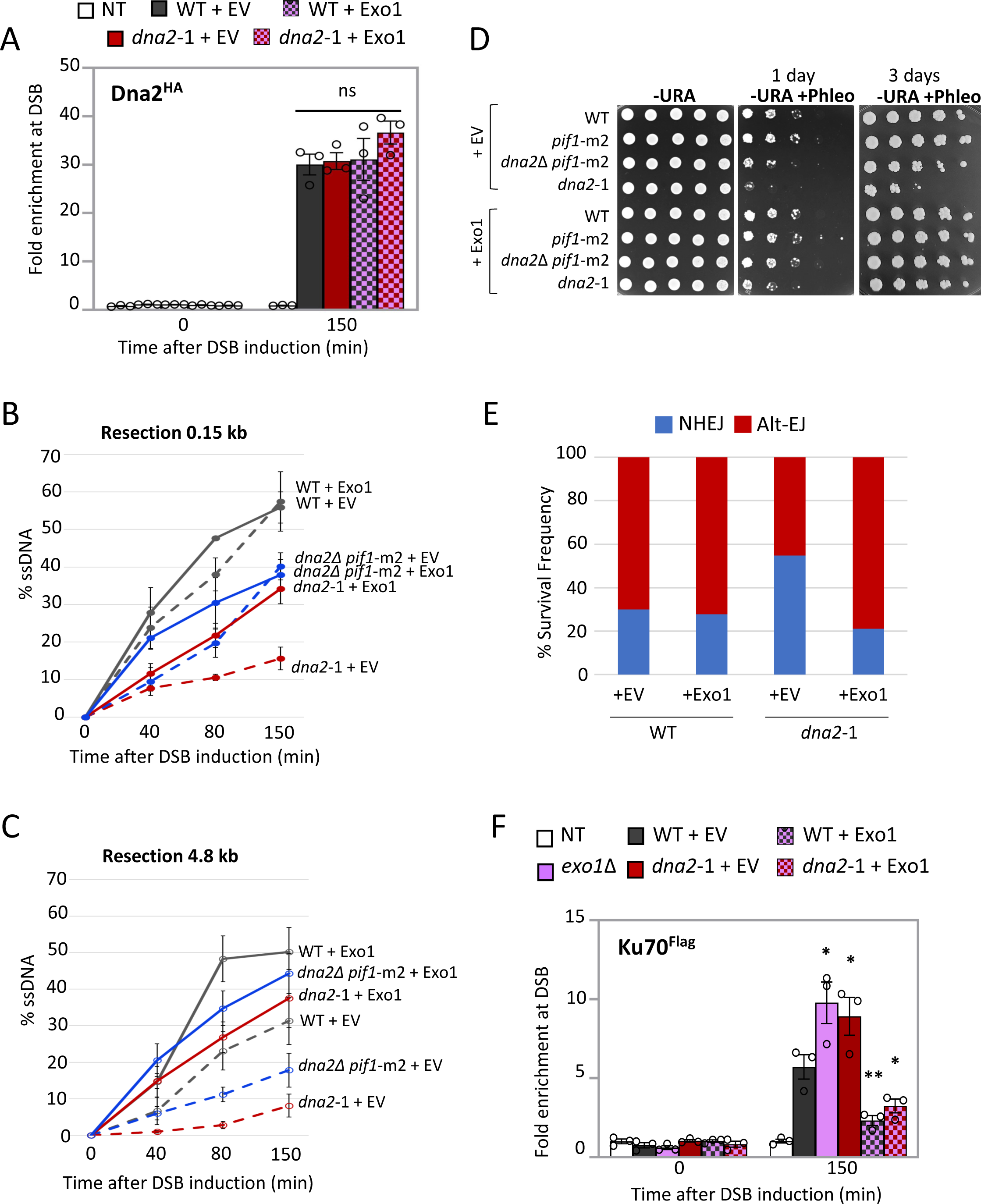
Overexpression of Exo1 restores the resection defect in *dna2*-1 cells. **(A)** Enrichment of Dna2^HA^ at 0.15kb from DSB, 0 min (no DSB induction) and 150 min after DSB induction in wild type (JC-4117), *dna2*-1 (JC-5707) with 2-micron empty vector (pRS426) or 2-micron plasmid encoding Exo1 (pEM-EXO1) and no tag control (JC-727). The fold enrichment is normalized to recovery at the *PRE1* locus. **(B-C)** qPCR-based resection assay of DNA 0.15kb and 4.8kb away from the HO DSB, as measured by % ssDNA, at 0, 40, 80 and 150 min post DSB induction in cycling cells in wild type (JC-727), *dna2*Δ *pif1*-m2 (JC-6005), and *dna2*-1 (JC-6007) with either 2-micron empty vector (pRS426) or 2-micron plasmid encoding Exo1 (pEM-EXO1). **(D)** Five-fold serial dilutions of wild type (JC-727), *dna2*Δ *pif1*-m2 (JC-6005), *pif1*-m2 (JC-6006) and *dna2*-1 (JC-6007) with + pEM-EXO1 or empty vector. Cells were spotted on -URA plate +/-3.0 μg/ml phleomycin and allowed to grow for 1-3 days. **(E)** Frequency ratio of NHEJ (blue) and alt-EJ/MMEJ (red) in wild type (JC-5903) and *dna2*-1 (JC-6105) with either the 2-micron plasmid encoding Exo1 (pRS425-EXO1) or the empty vector (pRS425) ctrl. **(F)** Enrichment of Ku70^Flag^ at 0.1 5kb from DSB, 0 min (no DSB induction) and 150 mins after DSB induction in wild type (JC-3964), *dna2*-1 (JC-6237) with either 2-micron empty vector (pRS426) or 2-micron plasmid encoding Exo1 (pEM-EXO1), *exo1*Δ (JC-6242) and no tag control (JC-727) was determined. The fold enrichment is normalized to recovery at the *PRE1* locus. For all the experiments -error bars represent the standard error of three replicates. Significance was determined using a 1-tailed, unpaired Student’s t test. All strains compared are marked (P<0.05*; P<0.01**) and compared to WT.

Lastly, we wanted to determine whether ectopic expression of EXO1 in *dna2*-1 mutants could restore the balance of NHEJ and alt-EJ/ MMEJ end joining repair. Again, we utilized the reporter system where cells were transformed with either pRS425-EXO1 for ectopic expression of Exo1 from a 2μ LEU2 plasmid or empty vector (EV) and where NHEJ and alt-EJ/ MMEJ were distinguished by growth on -URA media (Fig. S2F). The relative frequency of end-joining by alt-EJ/MMEJ and NHEJ in *dna2*-1 mutants was reversed upon expression of EXO1 (Figs. 2F and 4E). These findings are consistent with short-range resection being important for alt-EJ/MMEJ, but also with decreased Ku (Fig. 4F). In wild type cells, Exo1 expression also decreased the level of Ku recovered at the DSB, however the frequency of alt-EJ/MMEJ to NHEJ repair did not change, nor did short-range resection (Figs 4E, F).

## Discussion

We set out to understand why *dna2*-1 nuclease dead mutants were more sensitive to DSB causing agents than *dna2*Δ mutants. Our results showed that the presence of nuclease dead Dna2 at DSBs, localization of Exo1 decreased and inhibited resection more than disrupting either nuclease by deletion of *DNA2* or *EXO1*. The negative effect of *dna2*-1 at DSB was caused by Ku dependent inhibition of Exo1 localization (Fig. 5A, C). The *dna2*-1 phenotype was largely overcome by either deleting *KU70* or expressing Exo1, as both rescued the resection defect and phleomycin sensitivity. By contrast, neither Exo1 or Ku70 recovery changed in *dna2*Δ *pif1*-m2, however Nej1 levels markedly increased (Fig. 5B). Our work corroborates previous SGA screening where *dna2*-1 in combination with *exo1*Δ was viable [32], and suggests that the SL resulting from deletion of both *DNA2* and *EXO1* stems from something other than a combined loss of nuclease activity, unless *dna2*-1 has activity *in vivo* below the level of detection.

**Figure 5.**
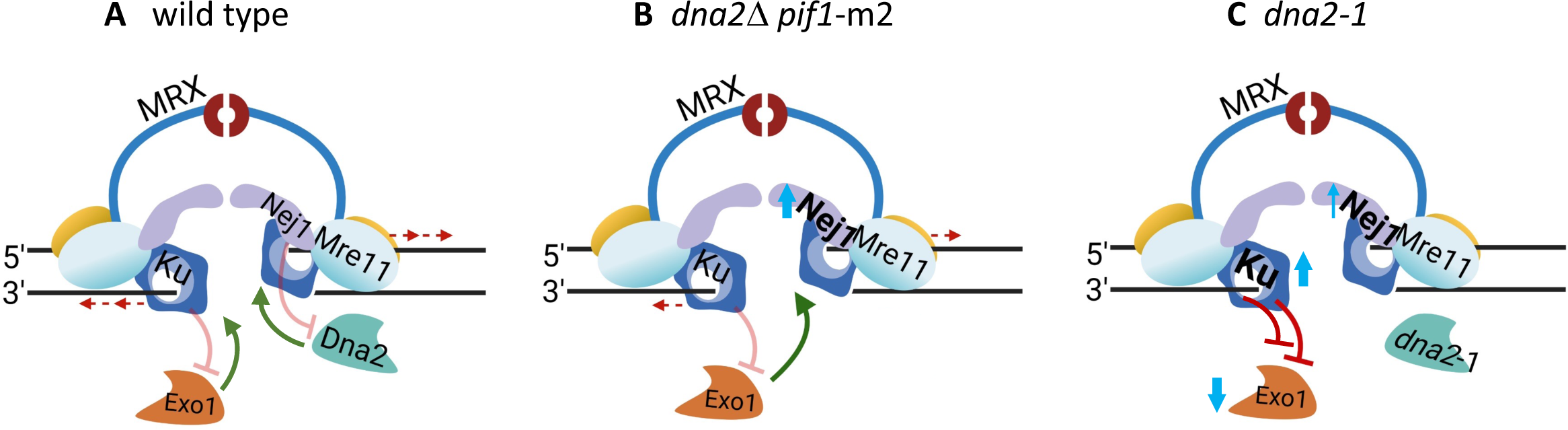
Model depicting how dna2 mutants impact DSB repair, where the presence of nuclease dead Dna2 at the break inhibits Exo1 nuclease through increased Ku. **(A)** The schematic shows the antagonistic relationship between Ku70/80 (Ku) and Exo1 and Nej1 and Dna2. Ku binding to DNA ends at the break, inhibiting the access of Exo1 nuclease to perform 5’ resection. Nej1 is a competitive inhibitor of Dna2, blocking interactions between Dna2 and its binding partners at DSBs [1, 5, 6, 8, 9, 15,16]. **(B)** Upon deletion of *DNA2,* Nej1 increased but Ku and Exo1 levels did not change. In *dna2*D *pif1*-m2 mutants, resection decreased ∼2-fold as Exo1 was the only functional long-range nuclease at the DSB, and the frequency of alt-EJ/MMEJ markedly increased. **(C)** In *dna2*-1 mutants, nuclease-dead Dna2 was recruited to the break site. Under this condition, there was a minor increase in Nej1. However, Ku increased, which in turn resulted in Exo1 inhibition. Therefore, in this mutant background, the functionality of both nucleases was compromised, resection was abrogated, and the frequency of NHEJ increased. The *dna2*-1 resection defect and phleomycin sensitivity were largely reversible either by ectopic expression of Exo1 or by deletion of Ku70. However, only Exo1 expression restored the balance of NHEJ and alt-EJ/MMEJ frequencies to levels observed in wild type cells.

The loss of Dna2 nuclease activity by a point mutation had a profound impact on the pathway used in DSB repair, and 5’ resection was central to the whole process. Ku and Nej1 are negative regulators of resection, and previous work by our lab and others showed there to be a division of labor for nuclease inhibition by these NHEJ factors [5, 6, 8, 9, 15, 16]. Nej1 interacts with the binding partners of Dna2, including Mre11, Sae2, and Sgs1 to inhibit its activity. However, Nej1 does not inhibit Exo1 recruitment. By contrast, Ku binding at the DSB directly inhibits the accessibility of Exo1 nuclease to DNA ends, and its recovery at the break site, whereas Dna2 can initiate resection in the presence of Ku [24, 25]. Thus, the antagonistic relationship between Nej1 and Dna2 is distinct, and independent, from the antagonistic relationship between Ku and Exo1 at DSB. The data presented here highlight a layer of complexity not previously reported in earlier work, where Dna2 fidelity impacts the balance of Ku and Exo1 at the break, a relationship paramount for ensuring DSBs are repaired through the least mutagenic pathway available.

The *dna2* mutants impacted end-joining differently in the reporter cell line where HR was not an option, a scenario relevant in G1 of the cell cycle. While NHEJ is error prone with the formation of small insertions and deletions, a more harmful outcome arises when resection initiates and alt-EJ/MMEJ is used, as it is more mutagenic. In *dna2*-1 mutants, the relative frequency of NHEJ increased and alt-EJ/MMEJ decreased, which was consistent with more Ku recovered at the break. The presence of Ku impacted the type of end-joining repair in *dna2*-1 more so than the pronounced defect to resection as both *dna2*-1 *exo1*Δ and *dna2*-1 *exo1*Δ ku70Δ showed similar resection defects, but NHEJ was Ku dependent (Fig. S3C). On the contrary, there was marked increase in alt-EJ/MMEJ in *dna2*Δ *pif1*-m2 mutants. This might have something to do with the fact that resection proceeded, although at reduced rate, in cells where *DNA2* was deleted. However, resection also equally decreased when *EXO1* was deleted and repair occurred predominately through NHEJ (Figs. 3B and S3C). In all, these data might provide information underscoring the cause of SL resulting from deletion of EXO1 and DNA2, as HR and both end-joining pathways would be disrupted.

The end-joining observations also suggest that the presence of Ku was not the only regulatory factor determining the type of end-joining repair. Rather our data support a model where the frequency of NHEJ to alt-EJ/MMEJ was impacted by the relative level of Ku in relation to other repair factors, namely Exo1 and Nej1. Exo1 promoted alt-EJ/MMEJ in the presence of Ku in *dna2* mutants, and we observed this under two experimental conditions. First, alt-EJ/MMEJ in *dna2*-1 mutants increased upon Exo1 expression (Fig. 4E), and secondly, alt-EJ/MMEJ increased in *dna2*Δ *pif1*-m2 mutants, where the levels of Exo1 and Ku70 were unaltered, but the level of Nej1 increased. (Figs. 2A, B and S3C). These phenotypic differences between *dna2*Δ and *dna2*-1 indicate the possibility of increased Nej1-dependent alt-EJ/MMEJ repair in *dna2*Δ cells [33, 34]. Further work is needed to elucidate whether Nej1 has a role in regulating end-joining; however, in support of this model, we recently reported that alt-EJ/MMEJ increased in aging cells as Ku declined, and Nej1 persisted DSBs [35].

Taken together, our data point out the dynamic interplay between Dna2 and Exo1 in DSB repair pathway choice and broaden the understanding of nuclease localization vs nuclease activity in DNA processing at break sites. The characterization of the two *dna2* mutants demonstrated that loss of DNA2 nuclease activity through different mutations resulted in different repair outcomes. The work has health relevance, and although *DNA2* deletion is embryonic lethal, point mutations are observed in diseases like Seckel syndrome and various kinds of cancer, with the corresponding P504→S mutation of *dna2*-1 in yeast seen in human cancers [28, 29].

### Experimental procedures

#### Media details

All the yeast strains used in this study are listed in Table S1 and were obtained by crosses. The strains were grown on various media in experiments described below. For HO-induction of a DSB, YPLG media is used (1% yeast extract, 2% bactopeptone, 2% lactic acid, 3% glycerol and 0.05% glucose). For the continuous DSB assay, YPA plates are used (1% yeast extract, 2% bacto peptone, 0.0025% adenine) supplemented with either 2% glucose or 2% galactose. For the mating type assays, YPAD plates are used (1% yeast extract, 2% bacto peptone, 0.0025% adenine, 2% dextrose).

#### Chromatin Immunoprecipitation

ChIP assays were performed as previously described [5]. Cells were cultured overnight in YPLG at 25°C. Cells were then diluted to equal levels (5 x 10^6^ cells/ml) and were cultured to one doubling (3-4 hrs) at 30°C. 2% GAL was added to the YPLG and cells were harvested and crosslinked at various time points using 3.7% formaldehyde solution. Cut efficiencies for all strains are shown in Table S2. Following crosslinking, the cells were washed with ice cold PBS and the pellet stored at -80°C. The pellet was re-suspended in lysis buffer (50mM Hepes pH 7.5, 1mM EDTA, 80mM NaCl, 1% Triton, 1mM PMSF and protease inhibitor cocktail) and cells were lysed using Zirconia beads and a bead beater. Chromatin fractionation was performed to enhance the chromatin bound nuclear fraction by spinning the cell lysate at 13,200 rpm for 15 minutes and discarding the supernatant. The pellet was re-suspended in lysis buffer and sonicated to yield DNA fragments (∼500bps in length). The sonicated lysate was then incubated αHA-, αFlag- or αMyc-antibody conjugated beads or unconjugated beads (control) for 2 hours at 4°C. The beads were washed using wash buffer (100mM Tris pH 8, 250mM LiCl, 150mM (αHA and αFlag) or 500mM (αMyc) NaCl, 0.5% NP-40, 1mM EDTA, 1mM PMSF and protease inhibitor cocktail) and protein-DNA complex was eluted by reverse crosslinking using 1% SDS in TE buffer, followed by proteinase K treatment and DNA isolation via phenol-chloroform-isoamyl alcohol extraction. Quantitative PCR was performed using the Applied Biosystem QuantStudio 6 Pro machine. PowerUp SYBR Green Master Mix was used to visualize enrichment at *MAT1* (0.15kb from DSB) and *PRE1* was used as an internal control (Table S2). HO cutting was measured in strains used to perform ChIP in Table S3.

#### Continuous DSB assay and identification of mutations in survivors

Cells were grown overnight in YPLG media at 25°C to saturation. Cells were collected by centrifugation at 2500rpm for 3 minutes and pellets were washed 1× in ddH_2_O and re-suspended in ddH_2_O. Cells were counted and spread on YPA plates supplemented with either 2% GLU or 2% GAL. 1×10^3^ total cells were plated on Glucose and 1×10^5^ total cells were plated on galactose. The cells were incubated for 3-4 days at room temperature and colonies were then counted on each plate. Survival was determined by normalizing the number of surviving colonies on the GAL plates to number of colonies on the GLU plates. 100 survivors from each strain were scored for the mating type assay as previously described [16] and at least 100 survivors were used to make a master-plate which was later replica-plated on -URA plates and the number of survivors on -URA plates were counted to determine the ratio of NHEJ and alt-EJ repair frequencies.

#### qPCR based Ligation Assay

As described previously [9], cells from each strain were grown overnight in 15ml YPLG to reach an exponentially growing culture of 1×10^7^ cells/ml. Next, 2.5ml of the cells were pelleted as ‘No break’ sample, and 2% GAL was added to the remaining cells, to induce a DSB. 2.5ml of cells were pelleted after a 3-hour incubation as timepoint 0 sample. After that, GAL was washed off and the cells were released in YPAD and respective timepoint samples were collected. Genomic DNA was purified using standard genomic preparation method by isopropanol precipitation and ethanol washing, and DNA was re-suspended in 100μL ddH_2_O. Quantitative PCR was performed using the Applied Biosystem QuantStudio 6 Flex machine. PowerUp SYBR Green Master Mix was used to quantify resection at HO6 (at DSB) locus. The PRE1 locus was used as an internal gene control for normalization. Signals from the HO6/PRE1 timepoints were normalized to ‘No break’ signals and % Ligation was determined. The primer sequences are listed in Table S2.

#### qPCR based Resection Assay

Cells from each strain were grown overnight in 15ml YPLG to reach an exponentially growing culture of 1×10^7^ cells/ml. Next, 2.5ml of the cells were pelleted as timepoint 0 sample, and 2% GAL was added to the remaining cells, to induce a DSB. Following that, respective timepoint samples were collected. Genomic DNA was purified using standard genomic preparation method by isopropanol precipitation and ethanol washing, and DNA was re-suspended in 100mL ddH_2_O. Genomic DNA was treated with 0.005μg/μL RNase A for 45min at 37°C. 2μL of DNA was added to tubes containing CutSmart buffer with or without inclusion of the *Rsa*I restriction enzyme and incubated at 37°C for 2hrs. Quantitative PCR was performed using the Applied Biosystem QuantStudio 6 Flex machine. PowerUp SYBR Green Master Mix was used to quantify resection at the RsaI cut site 0.15Kb DSB (in the *MAT1 locus*) and 4.8 Kb. PRE1 was used as a negative control and the primer sequences are listed in Table S2. *Rsa*I cut DNA was normalized to uncut DNA as previously described to quantify the %ssDNA [30]. HO cutting was measured in strains for resection Table S4.

## Supporting information

This article contains supporting information.

## Acknowledgments

We thank Lorraine Symington for generously providing us with the plasmid overexpressing EXO1, pEM-EXO1.

## Funding and additional information

This work was supported by operating grants from CIHR MOP-82736; MOP-137062 and NSERC 418122 awarded to J.A.C. The authors declare that they have no conflicts of interest with the contents of this article.

## Conflict of interest

The authors declare that they have no conflict of interest with the contents of this article.

## Supporting Information

**Figure S1.**
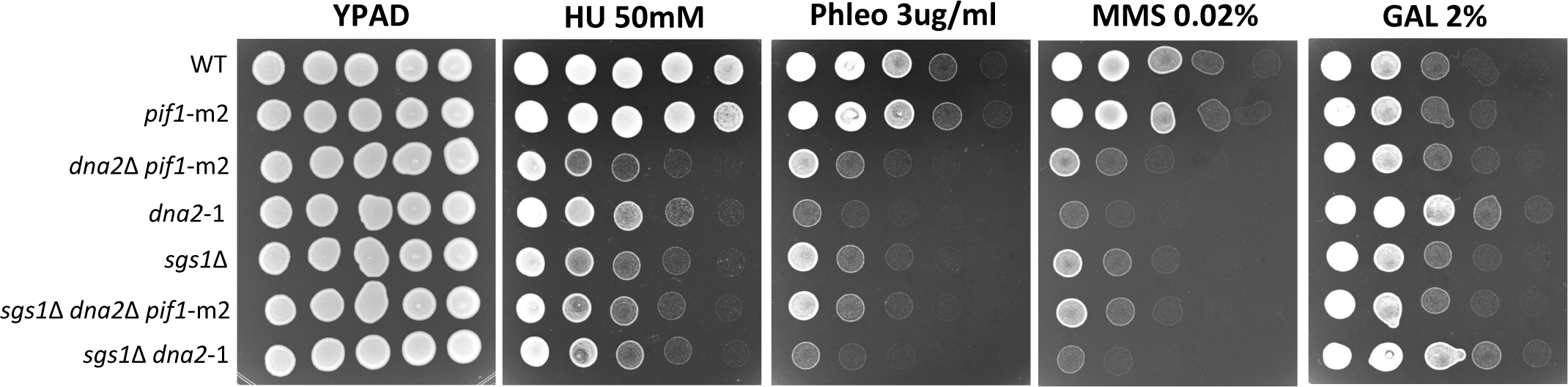
Sensitivity of various *dna2* and *sgs1* mutants to genotoxic stress. Five-fold serial dilutions of wild type (JC-727), *dna2*Δ *pif1*-m2 (JC-6005), *pif1*-m2 (JC-6006), *dna2*-1 (JC-6007), *sgs1*Δ (JC-3757), *sgs1*Δ *dna2*Δ *pif1*-m2 (JC-6101) and *sgs1*Δ *dna2*-1 (JC-5745) were spotted on YPAD, 50mM HU, 3.0 μg/ml phleomycin, 0.02% MMS and 2% GAL containing plates.

**Figure S2.**
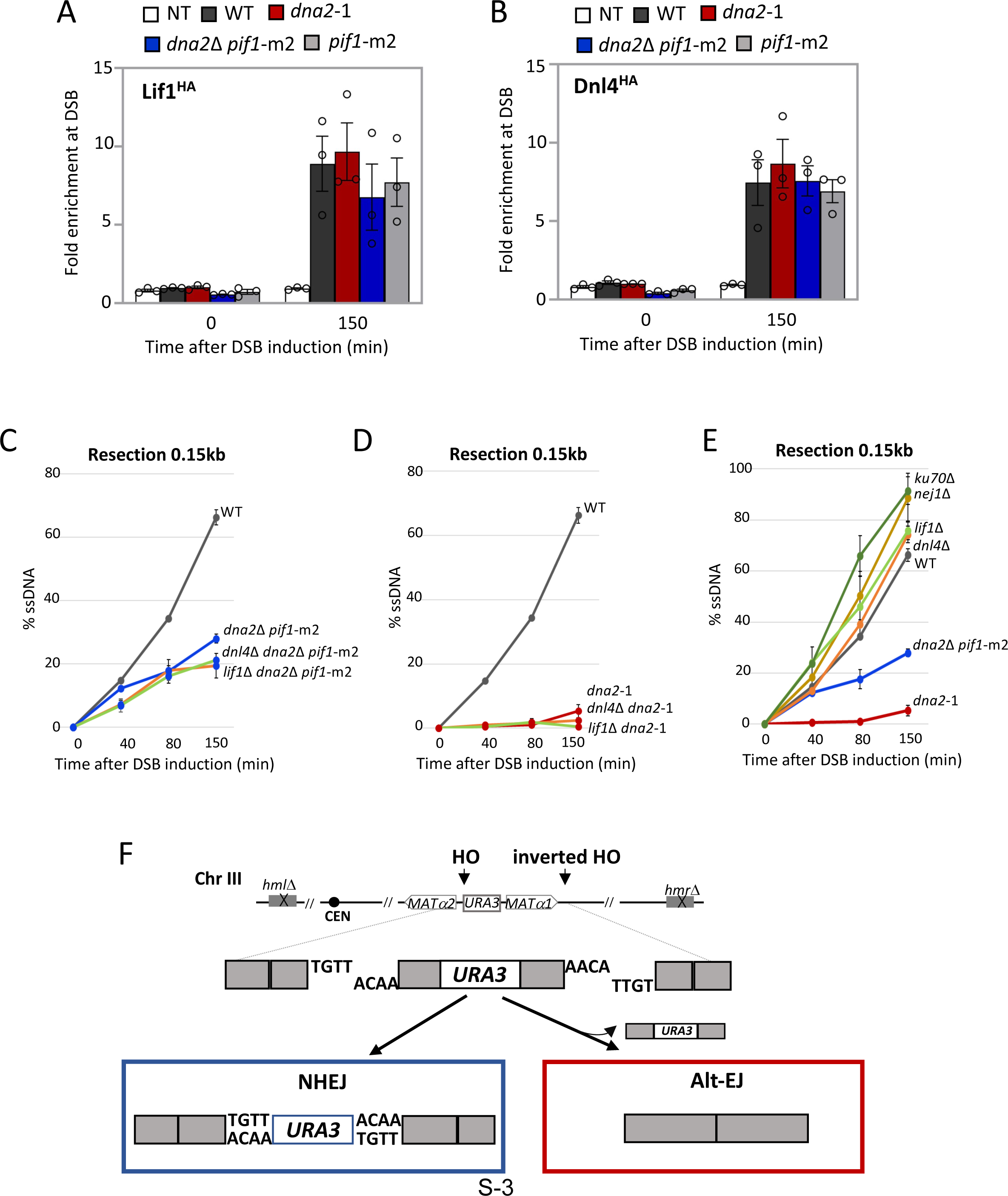
End-joining factors at DSBs in the dna2 mutants. **(A)** Enrichment of Lif1^HA^ at 0.15kb from DSB, 0 min (no DSB induction) and 150 min after DSB induction in wild type (JC-3319), *dna2*-1 (JC-5834), *dna2*Δ *pif1*-m2 (JC-6110), *pif1*-m2 (JC-6136) and no tag control (JC-727). **(B)** Enrichment of Dnl4^HA^ at 0.15kb from DSB, 0 min (no DSB induction) and 150 min after DSB induction in wild type (JC-5672), *dna2*-1 (JC-5843), *dna2*Δ *pif1*-m2 (JC-6123), *pif1*-m2 (JC-6135) and no tag control (JC-727). **(C-E)** qPCR based resection assay of DNA 0.15kb away from the HO DSB, as measured by % ssDNA, at 0, 40, 80 and 150 min post DSB induction in cycling cells in wild type (JC-727), *dna2*Δ *pif1*-m2 (JC-6005), *dna2*-1 (JC-6007), *ku70*Δ (JC-1904), *nej1*Δ (JC-1342), *lif1*Δ (JC-1343), *dnl4*Δ (JC-3290), *lif1*Δ *dna2*Δ *pif1*-m2 (JC-6121), *lif1*Δ *dna2*-1 (JC-5890), *dnl4*Δ *dna2*Δ *pif1*-m2 (JC-6119) and *dnl4*Δ *dna2*-1 (JC-5800). **(F)** Schematic representation of regions around the two HO cut sites on chromosome III. Following HO endonuclease break induction, NHEJ repair joins the complementary ends of the *URA3* gene with the other ends of the break site. From two simultaneous cuts, noncomplementary ends are generated, in repair progresses through alt-EJ/MMEJ repair the *URA3* gene is deleted.

**Figure S3.**
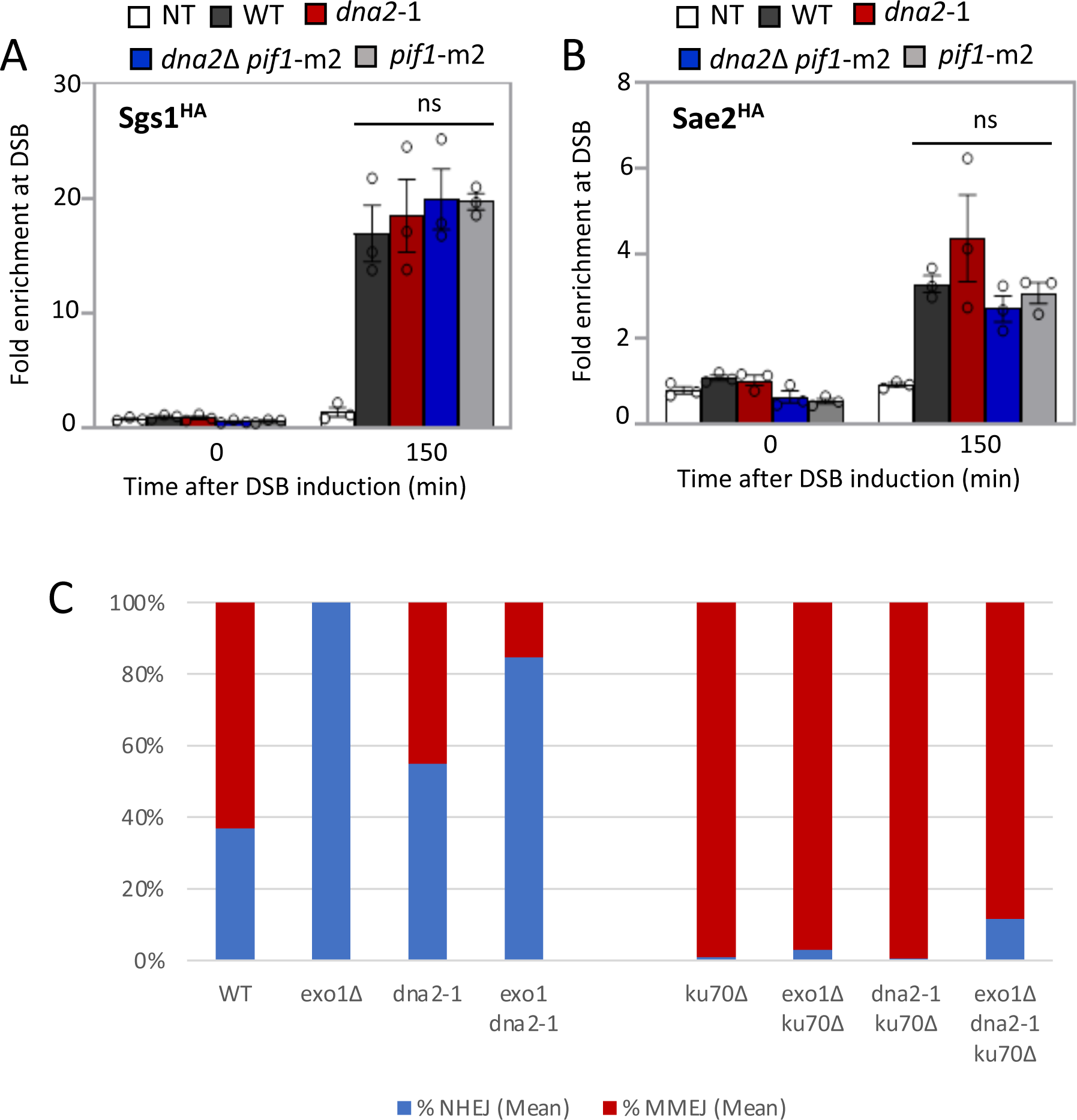
HR factors at DSBs in *dna2* mutants. **(A)** Enrichment of Sgs1^HA^ at 0.15kb from DSB, 0 min (no DSB induction) and 150 min after DSB induction in wild type (JC-4135), *dna2*-1 (JC-5681), *dna2*Δ *pif1*-m2 (JC-6112), *pif1*-m2 (JC-6114) and no tag control (JC-727). **(B)** Enrichment of Sae2^HA^ at 0.15kb from DSB, 0 min (no DSB induction) and 150 min after DSB induction in wild type (JC-5116), *dna2*-1 (JC-5682), *dna2*Δ *pif1*-m2 (JC-6108), *pif1*-m2 (JC-6109) and no tag control (JC-727). **(C)** Survival frequencies depicting the ratio of NHEJ (blue) and alt-EJ (red) repair frequencies in wild type (JC-5903), *exo1*Δ (JC-6222), *dna2*-1 (JC-6105), *exo1*Δ *dna2*-1 (JC-6328), *ku70*Δ (JC-6195), *ku70*Δ *exo1*Δ (JC-6272), *ku70*Δ *dna2*-1 (JC-6273) and *ku70*Δ *exo1*Δ *dna2*-1 (JC-6326).

**Figure S4.**
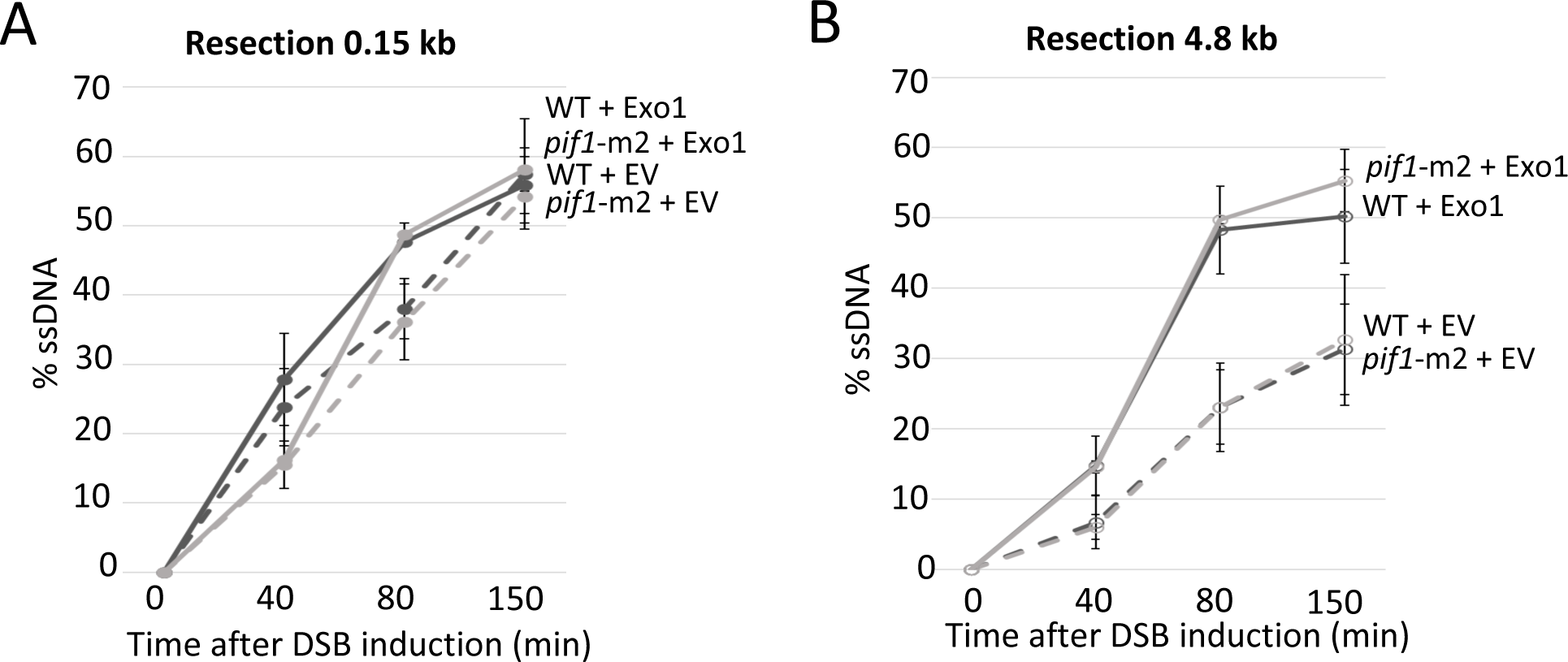
Resection in *pif1*-m2 mutants is unaltered by Exo1 expression. **(A-B)** qPC-based resection assay of DNA at two distances, 0.15kb and 4.8kb, from the HO DSB as measured by % ssDNA, at 0, 40, 80 and 150 mins. after DSB induction in cycling cells. Resection is compared in wild type (JC-727) *and pif1*-m2 (JC-6006) with 2-micron plasmid encoding Exo1 (pEM-EXO1) or empty vector (EV) [15].

**Table S1:**
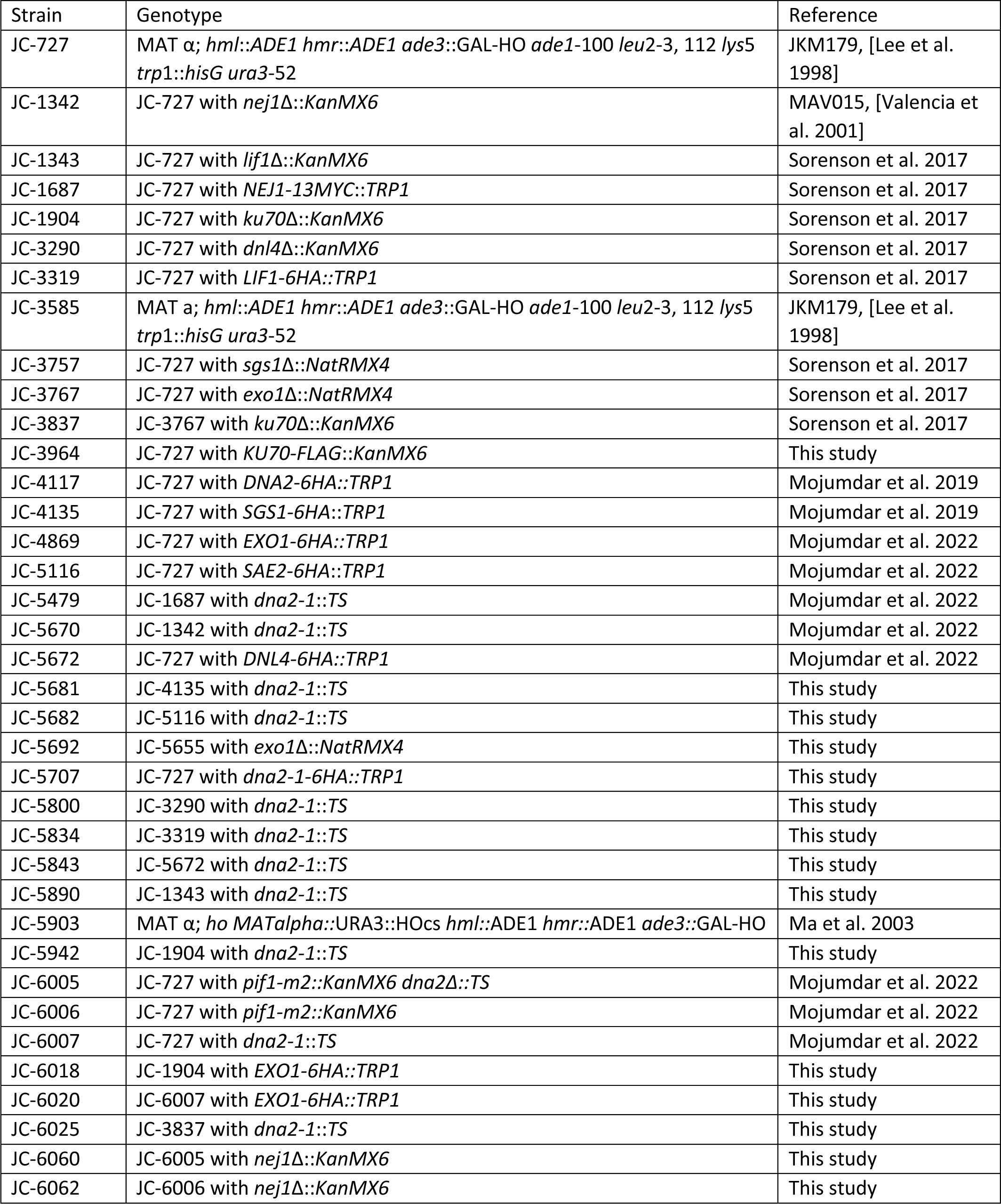
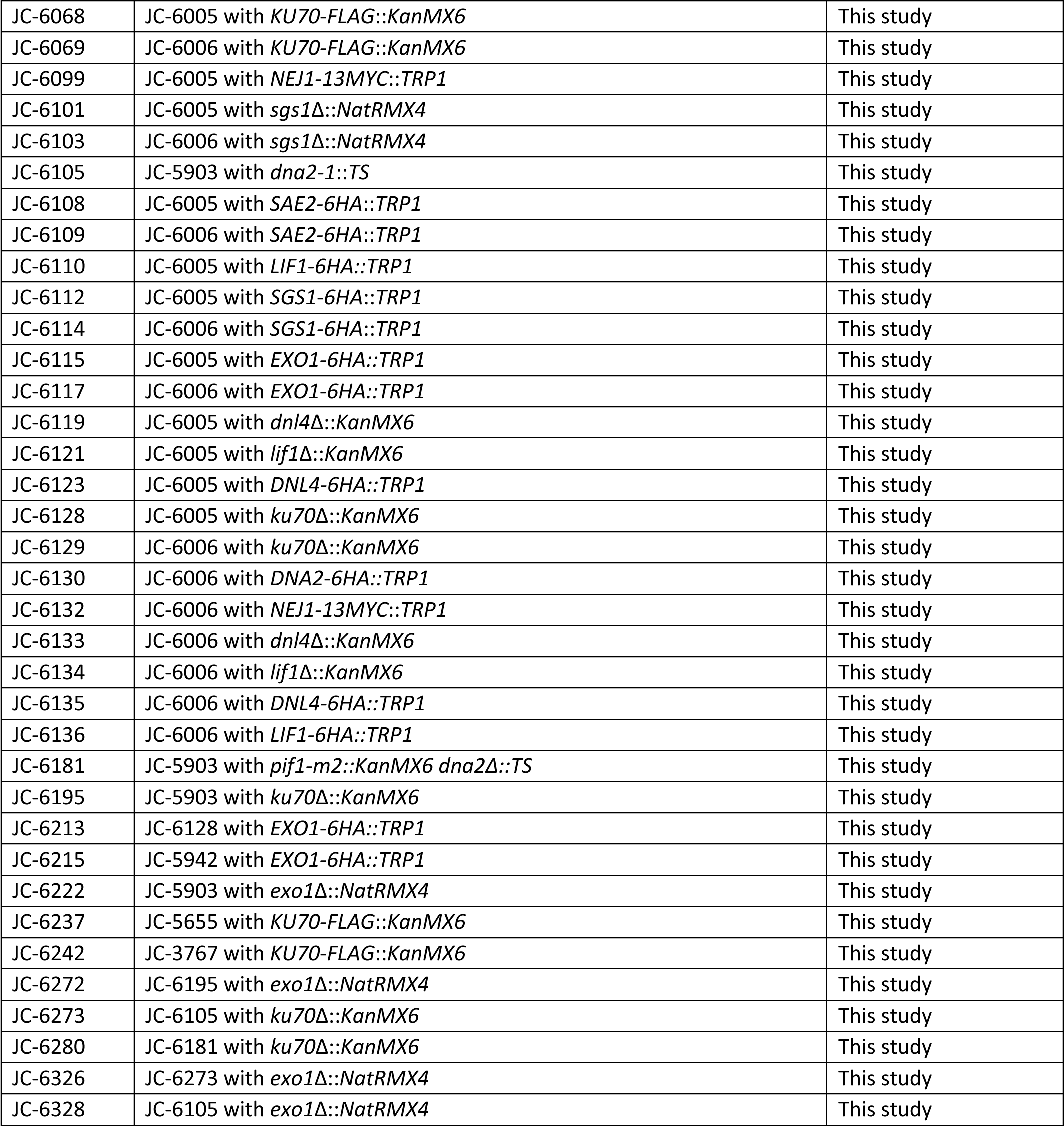
*S. cerevisiae* strains used in this study.

**Table S2:**
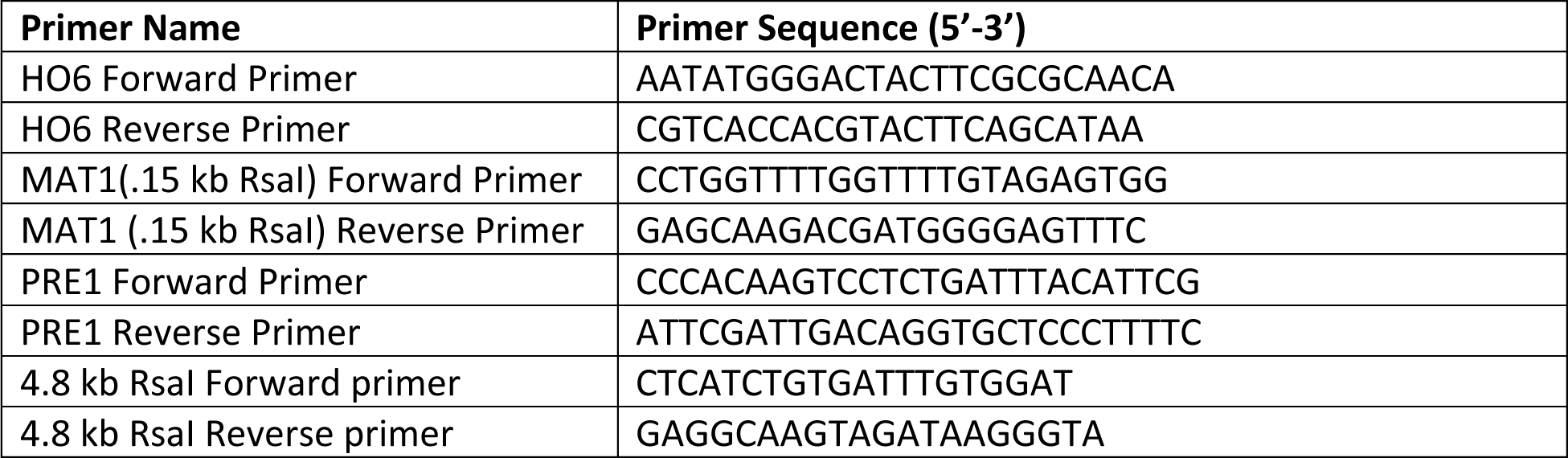
Primers used in these studies.

**Table S3:** DSB cut efficiency for strains used in ChIP.

| **Strain** | **DSB cut % at**<br>**150min** |
| --- | --- |
| JC-3964 | 95.65 |
| JC-4117 | 98.94 |
| JC-4869 | 99.27 |
| JC-5707 | 98.93 |
| JC-6018 | 98.36 |
| JC-6020 | 98.82 |
| JC-6068 | 92.63 |
| JC-6069 | 84.81 |
| JC-6115 | 84.58 |
| JC-6117 | 80.90 |
| JC-6130 | 95.65 |
| JC-6215 | 98.94 |
| JC-6237 | 99.27 |
| JC-6242 | 98.93 |

**Table S4:** DSB cut efficiency for strains used in resection assay.

| Strain | DSB cut (%) |  |  |
| --- | --- | --- | --- |
|  | 40min | 80min | 150min |
| JC-727 | 50.95 | 85.95 | 95.45 |
| JC-1904 | 89.11 | 88.98 | 97.66 |
| JC-3757 | 52.12 | 85.68 | 94.85 |
| JC-3767 | 51.67 | 77.16 | 91.32 |
| JC-3837 | 91.96 | 97.71 | 99.04 |
| JC-5372 | 75.22 | 88.84 | 93.10 |
| JC-5655 | 39.69 | 79.06 | 90.56 |
| JC-5669 | 33.13 | 79.71 | 83.59 |
| JC-5673 | 51.12 | 69.31 | 81.26 |
| JC-5692 | 50.54 | 65.28 | 81.45 |
| JC-5745 | 76.57 | 92.11 | 97.03 |
| JC-5748 | 46.41 | 79.62 | 91.65 |
| JC-5942 | 91.68 | 98.00 | 98.96 |
| JC-6005 | 94.39 | 99.13 | 99.09 |
| JC-6006 | 82.96 | 93.15 | 94.72 |
| JC-6025 | 78.37 | 93.97 | 94.31 |
| JC-6098 | 75.89 | 94.17 | 95.25 |
| JC-6101 | 76.68 | 84.78 | 94.48 |
| JC-6128 | 77.22 | 83.92 | 93.96 |
| JC-6152 | 84.61 | 89.88 | 90.49 |

## Notes

### Competing Interest Statement

The authors have declared no competing interest.

